# CFTR mutation leads to intrinsic dysfunction in neutrophils from people with Cystic Fibrosis

**DOI:** 10.1101/2025.06.08.656500

**Authors:** Frank H Robledo-Avila, Raul Rascon, Alejandra Montanez-Barragan, Veronica Loyo-Celis, Harpreet Singh, Karen S McCoy, Benjamin T Kopp, Santiago Partida-Sanchez

**Affiliations:** Center for Microbe and Immunity Research, Abigail Wexner Research Institute at Nationwide Children’s Hospital, Columbus, OH, USA; Department of Physiology and Cell Biology, College of Medicine, The Ohio State University, Columbus, OH, USA; Department of Molecular Cellular and Developmental Biology, The Ohio State University, Columbus, OH, U.S.A; Division of Pulmonary Medicine, Nationwide Children’s Hospital, Columbus, OH, USA; Department of Pediatrics, College of Medicine, The Ohio State University, Columbus, OH, USA; Division of Pulmonology, Asthma, Cystic Fibrosis, and Sleep, Emory University School of Medicine, Atlanta, GA, USA; Children’s Healthcare of Atlanta, Atlanta, GA, USA

**Keywords:** Cystic fibrosis, Neutrophils, CFTR modulators, NETs, CFTR

## Abstract

Cystic fibrosis (**CF**), a common genetic disease, is caused by a defective CF-transmembrane conductance regulator (**CFTR**). People with CF (**pwCF**) are prone to develop infections by opportunistic pathogens, including *Burkholderia cenocepacia*, leading to chronic inflammation and lung function loss. Neutrophils, the most abundant cells in the chronically inflamed lungs of pwCF, release granular proteins and oxidative products that contribute to tissue damage. The CFTR modulators are a new treatment for pwCF aiming to correct the subcellular location and function of the CFTR ion channel. The triple modulator combination of Elexacaftor, Tezacaftor, and Ivacaftor (**ETI**) or Trikafta^®^ has significantly improved clinical symptoms and overall provided a better quality of life for pwCF. The mechanism by which the CFTR modulators help to restore the antimicrobial functions of neutrophils is unknown. The present study demonstrates that neutrophils functionally express CFTR and reveals how ETI modifies subcellular CFTR trafficking in CF neutrophils. In addition, ETI treatment reduces intracellular chloride levels in human neutrophils, indicating activation of CFTR-dependent chloride efflux (outflow). Finally, ETI treatment also reestablished the intracellular antimicrobial killing of CF neutrophils by potentiating NADPH oxidase activity and producing Neutrophil Extracellular Traps (NETs). Together, our findings suggest that CFTR has an essential role in controlling neutrophil functions and that the CFTR modulators improve the health of pwCF by restoring the antimicrobial functions of CF neutrophils.

Graphical abstract.
**(A**) The F508del defective CFTR protein cannot reach the plasma membrane in CF neutrophils, which increases the intracellular concentrations of Cl^−^ ions, and allows other ions to be internalized, including Na^+^ and Ca^2+^. This ionic imbalance affects the NADPH oxidase, leading to a reduced preactivation response and consequently impacting NADPH oxidase-dependent antimicrobial mechanisms, including intracellular antimicrobial killing and NETosis. **(B)** Treating with ETI restores the CFTR expression in the plasma membrane of CF neutrophils, increasing the Cl^−^ efflux and regulating the intracellular levels of Na^+^ and Ca^2+^, leading to correcting the NADPH oxidase, which results in potentiating the intracellular antimicrobial killing and NETosis. Biorender^TM^ tools generated the images.

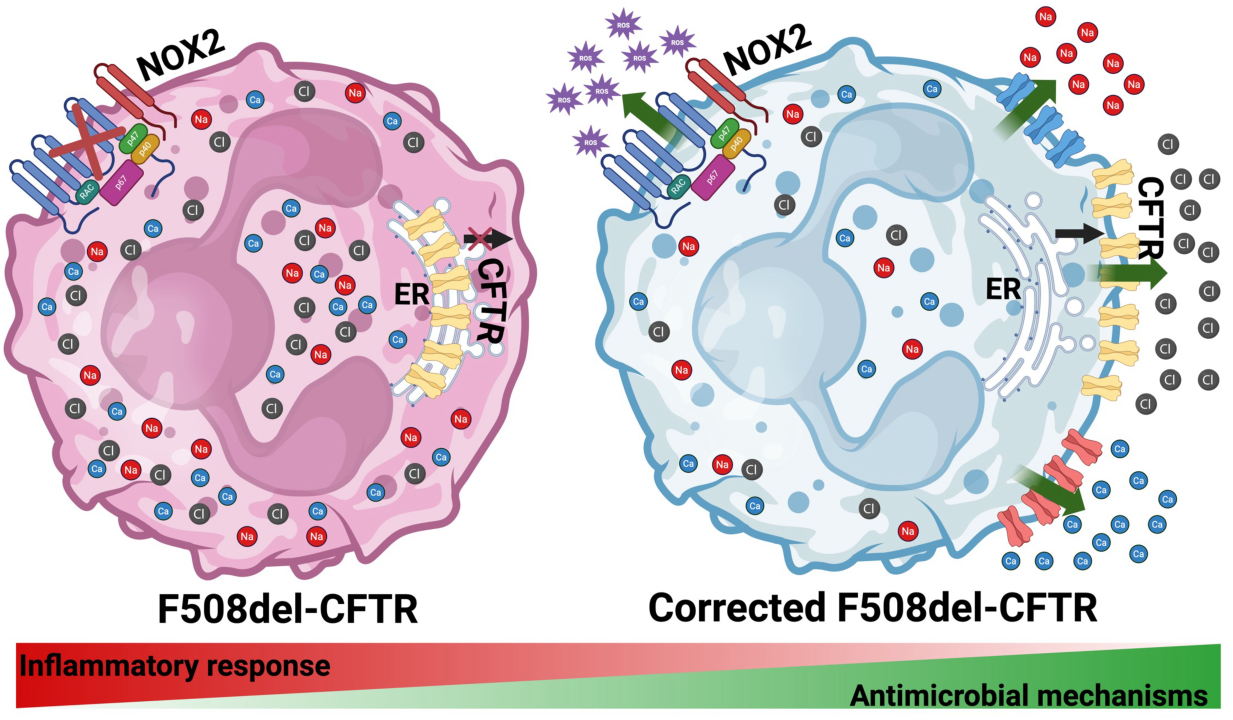

## INTRODUCTION

Cystic fibrosis (CF) is an autosomal recessive inherited disease that affects at least one hundred thousand people worldwide and remains with no cure [1]. Mutations in the gene located at the chromosome 7 position 31.2 (7q31.2) cause CF, leading to an abnormal or absent Cystic Fibrosis Transmembrane-Conductance Regulator (**CFTR**) protein [2–5]. CFTR is a transmembrane ATPase (ATP-binding cassette) that also works as a chloride (**Cl^−^**) and bicarbonate channel (HCO_3_^−^), regulated by cyclic adenosine monophosphate (**cAMP**) and protein kinase A phosphorylation [6, 7]. More than 2,100 mutations have been described to cause CF [8], with the F508del variant the most common in people with CF (**pwCF**). Deleting the phenylalanine residue at position 508 is present in one or both alleles in around 85% of pwCF, in which the CFTR protein is misfolded, and thereby, it gets trapped in the Endoplasmic Reticulum (**ER**) [9].

CF is a multisystemic disease. In the lungs, CFTR dysfunction causes abnormal transport of ions and dehydration of the epithelial surface, leading to accumulating thick mucus and establishing bacterial biofilms and chronic inflammation, resulting in tissue damage [5]. Most pwCF are prone to develop infections by opportunistic pathogens, including *Pseudomonas aeruginosa*, *Staphylococcus aureus* and *Burkholderia cenocepacia* [10–12]. Accumulating myeloid cells characterizes the lung inflammation upon infection in pwCF, with neutrophils being the most abundant cells in tissue [13]. Despite predominant neutrophilic inflammation in CF-infected lungs, bacterial infection cannot be satisfactorily resolved. This unresolvability may be because the polymorphonuclear cells (**PMNs**) demonstrate intrinsic antimicrobial defects, including a reduced oxidative burst, an excessive primary degranulation [14], delayed Neutrophil Extracellular Traps (NETs) production, and reduced intracellular antimicrobial killing against CF pathogens [15].

CFTR modulator therapies have been introduced to correct some functional defects in the CFTR ion channel [16, 17]. Recently, the triple combination, Elexacaftor/Tezacaftor/Ivacaftor (**ETI**) or Trikafta^®^, is one of the most widely used CFTR modulator therapies [5]. Clinical symptoms of pwCF under ETI treatment exhibit improvement in Forced Expiratory Volume at one second (FEV_1_), cough reduction, nutrition intake, and reduction in bacterial burden [18–20]. ETI treatment in vitro has demonstrated effects on CF airway epithelial cells with restored CFTR activity and increased rate of wound healing [21–23]. Additionally, our group has proven that treating with ETI restored CFTR expression in the plasma membrane and re-established CFTR-dependent current and Cl^−^ efflux in CF macrophages [24].

Neutrophils are the most abundant leukocytes in the lungs of pwCF during inflammation, and their granular proteins and oxidative burst promote profuse damage in the lungs of pwCF [25]. Previous report indicated some ETI effects on neutrophilic inflammation in pwCF, including reducing the number of neutrophils and granular markers in the sputum of pwCF at least 3 months post-treatment with ETI [26]. Another study showed no differences in total neutrophil and monocyte counts in blood and neutrophilic activation markers in pwCF after treatment with ETI as compared to healthy donors [27]. However, the effect of ETI on neutrophils at the cellular and molecular level has yet to be explored. In this work, we use in vitro and ex vivo approaches to establish functional expression of the Cl^−^ channel CFTR in neutrophils. The data demonstrate that treating with ETI can significantly improve neutrophil dysfunction in F508del neutrophils. Enhanced CFTR-dependent currents demonstrated the functional CFTR protein expression in neutrophils, which resulted in lower intracellular Cl^−^ levels in CF-treated neutrophils. Additionally, CFTR modulators markedly improve the defective antimicrobial killing properties of CF neutrophils, both in vitro and in vivo. The proposed mechanism of ETI in neutrophils is by restoring CFTR Cl^−^ channel functionality at the plasma membrane and activating the NADPH oxidase-dependent pathway, which promotes the release of NETs and restores effective antimicrobial killing of neutrophils.

## MATERIALS AND METHODS

### Bacterial culture

*Burkholderia cenocepacia* (*B. cenocepacia*) strain K56-2 batch was stored at -80 °C. One day before the experiment, bacteria were thawed from a frozen vial and streaked on *Burkholderia cepacia* selective agar plates (Thermofisher Scientific) and cultured at 37 °C overnight. The next day, a colony was picked and cultured in Luria-Bertani broth (Difco, MD) at 37 °C with shaking. Bacteria concentrations were adjusted based on absorbance at 600 nm.

### Isolation of human blood polymorphonuclear cells

Human donors gave written informed consent for blood donation as approved by the Institutional Review Board of Nationwide Children’s Hospital (Columbus, OH, USA; IRB16-01020). Written consent from legal guardians of minors was obtained with written assent from minors (9 to 17 years). Inclusion criteria included pwCF with at least one copy of the F508del CFTR variant (**Table 1**). Exclusion criteria included history of *B. cepacia* complex culture positivity, chronic immunosuppression, and history of transplantation. Blood was collected (20 mL per subject) in heparinized tubes (BD, Franklin Lakes, NJ). Negative selection (Stemcell Technologies, Vancouver, BC, Canada) purified the human peripheral blood neutrophils according to the manufacturer’s directions.

**Table 1.**
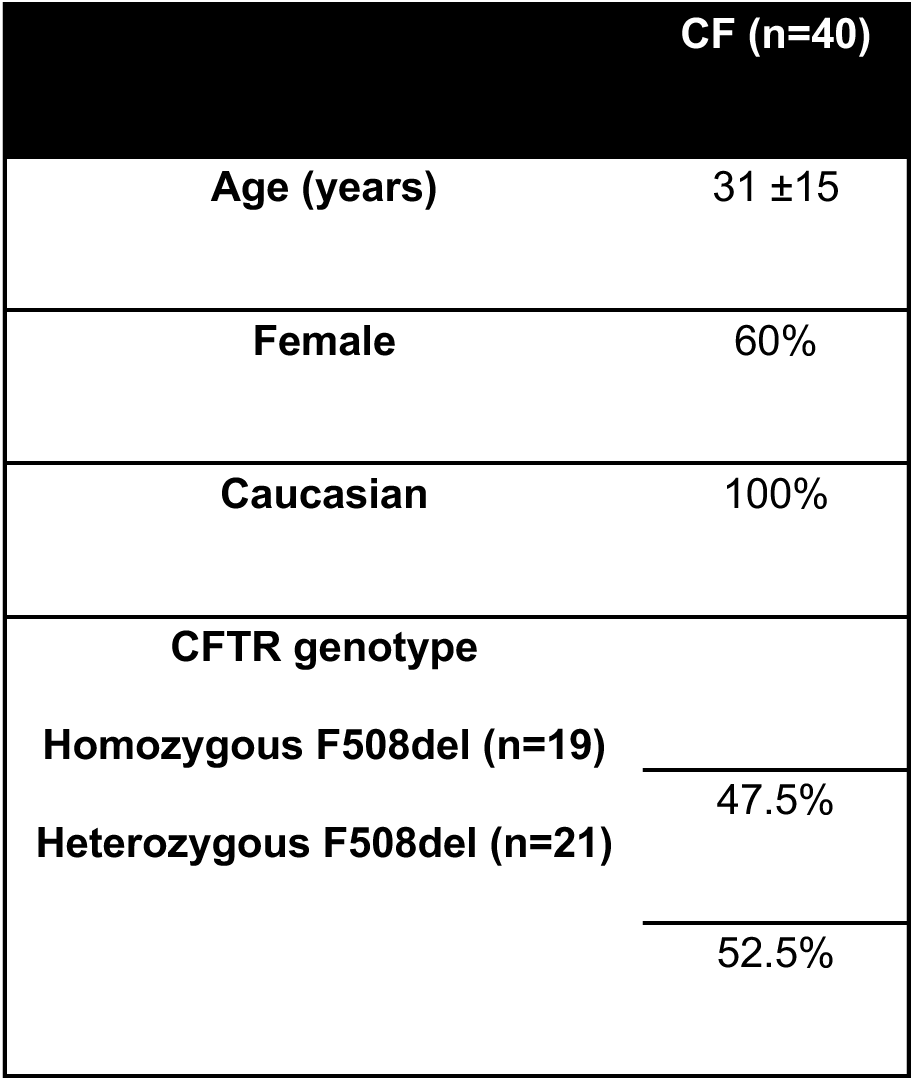
Participants demographics.

### Quantification of bacterial colony-forming units

Non-CF (Healthy donors) or CF neutrophils (10^6^ cells) were treated with Elexacaftor/Tezacaftor/Ivacaftor (**ETI**) (5 μM/5 μM/ 1μM) for one hour at 37 °C, then neutrophils were infected with *B. cenocepacia* with a Multiplicity of Infection of 10 (**MOI** = 10) in Roswell Park Memorial Institute (RPMI) 1640 medium with 2% fetal bovine serum for 45 minutes. After washing and centrifuging the cells at 1000 rpm for 7 minutes twice, the supernatants were removed, and the pellets resuspended and seeded in 24-well plates, and then incubated for three hours at 37 °C. To quantify intracellular bacteria, lysed neutrophils [0.1% Triton X-100 (Sigma Aldrich, St. Louis, MO)] were serially diluted on LB agar plates to quantify the free bacteria. The relative percent of antimicrobial killing was calculated by dividing colonly-forming units (**CFU**) from non-CF by CFU from CF neutrophils and then multiplying by 100.

### Measurement of reactive oxygen species

Neutrophils (10^5^ cells) were seeded in 96-well plates, then 10^−4^ M luminol added, followed by stimulating with 5×10^−8^ M phorbol 12-myristate 13-acetate (**PMA**). The luminescence plate reader Synergy H1 multimode plate reader (Biotek, Winooski, VT) acquired the kinetics. Additionally, neutrophils were prepared as mentioned above, 30 minutes after stimulating with 5×10^−8^ M PMA. Neutrophils were stained with cellROX green (Thermofisher Scientific) according to the manufacturer’s instructions, and the Synergy H1 multimode plate reader scanned the plates at 488/521 nm.

To detect hypochlorite or highly reactive oxygen species (**hROS**), neutrophils were prepared as mentioned before, followed by stimulating with 5×10^−8^ M PMA and adding 10 nM of Diphenyleneiodonium chloride (DPI) or combining 5 μM Elexacaftor/5 μM Tezacaftor/1 μM Ivacaftor (ETI). Then 5 μM aminophenyl fluorescein was added to the neutrophils to detect hROS. The Synergy H1 multimode plate reader detected hypochlorite production at 488/521 nm.

### Western blot and Immunoprecipitation

For measuring phosphorylation of the subunit phos-p47^phox^, PMN were seeded in 6-well plates (4×10^6^ cells per well). Neutrophils were treated with ETI for one hour, followed by stimulating with 5×10^−8^ M PMA or *B. cenocepacia* (MOI = 10) for 15 and 30 minutes. Unstimulated cells under the same conditions served as controls. Cells were lysed in TN1 lysis buffer (50 mM Tris pH 8.0, 10 mM EDTA, 10 mM Na_4_P_2_O_7_, 10 mM NaF, 125 mM NaCl, 9 mM Na_3_VO_4,_ and 1% Triton-X100). TN1 buffer was supplemented with 3X Halt Protease Enzyme Cocktail (Invitrogen). Lysates were spun at 13000xg for 10 minutes at 4 °C and then heated for 5 minutes at 96 °C in 4X or 6X Laemmli Sample Buffer. Sodium dodecyl sulfate polyacrylamide gel electrophoresis (**SDS-PAGE**) was made in 10% Bis-Tris at 100 V, and gels were transferred to polyvinylidene difluoride membranes for 40 minutes at 14 V under Semi-Dry conditions (Bio-Rad). Transferred membranes were blocked in 5% BSA for one hour. After blocking, Abs directed against phosphorylated p47p^phox^ (Ser 370) or (Ser 345) (Cell Signaling) were diluted 1:1000 in 5% BSA, 0.1% TBST-Tween 20, and membranes incubated overnight. On the next day, membranes were washed for 6 minutes (5 times) and incubated one hour with HRP-linked anti-rabbit IgG 1:7500. Chemiluminescence was detected with Super Signal Femto ECL Kit in a ChemiDoc MR machine (BioRad). Finally, membranes were stripped using Invitrogen Stripping Buffer and Total P47 (Cell Signaling) and GAPDH or β-Actin were used as loading control.

For detecting CFTR, 10^7^ neutrophils were pelleted in 1.5 mL conical tubes and resuspended in 100 μL of PBS with 3X Halt Protease Enzyme Cocktail (Invitrogen). Immediately, 100 μL of boiling 6X sample buffer was added to the cells and the mixture was boiled for 7 minutes at 96°C. After cooling, cell lysates were separated in 4-16% SDS-PAGE at 100 V. The gel was transferred overnight in CAPS buffer containing 10% methanol at 5V. Transferred membranes were blocked in 5% BSA for one hour. After blocking, anti-CFTR clone TJA9 acquired from the CFTR antibody distribution program at The University of North Carolina, Chapel Hill (UNCCH), or clone 24.1 (R&D Biosystems) were diluted 1:1000 in 5% BSA, 0.1% TBST-Tween 20 and incubated overnight. Next day, membranes were washed for 6 minutes 5 times and incubated one hour with HRP linked anti mouse IgG 1:7500. Chemiluminescence was detected with Super Signal Femto ECL Kit in a ChemiDoc MR machine (BioRad). Finally, membranes were stripped using Invitrogen Stripping Buffer and GAPDH or β-Actin were used as loading control.

### CFTR-mediated chloride efflux

The protocol was followed as previously described [28]. Briefly, neutrophils were stained with 1 mM N-ethoxycarbonylmethyl-6-Methoxyquinolinium Bromide [(**MQAE**) Thermofisher] for 30 minutes at 37 °C in HBSS media with Ca^2+^ and Mg^2+^ (Gibco), then the cells were washed and adjusted to 10^6^ neutrophils per tube. Flow cytometry (LSR Fortessa, BD Biosciences) recorded the kinetics using the UV laser 355-379/28 BP. The basal fluorescence was recorded up to 60 seconds, followed by stimulating CFTR with 20 μM Forskolin (Sigma)/50 μM 3-isobutyl-1-methylxanthine (**IBMX**, Sigma) up to 600 seconds and then 20 μM CFTR inhibitor 172 (CFTR_inh-172_, Selleckchem) was added, the kinetics were recorded up to 900 seconds.

We measured CFTR-mediated halide efflux using the method previously described [29] with some modifications. Briefly, PMNs were stained with 1 mM MQAE in efflux solution (135 mM NaNO_3_, 1 mM CaSO4, 1 mM MgSO4, 2.4 mM K2HPO4, 0.6 mM KH2PO4, 10 mM HEPES, and 10 mM glucose) for 30 minutes at 37 °C, then cells were washed, and NaI buffer (135 mM NaI in efflux buffer) was added to the cells. Flow cytometry recorded cell properties as described above.

### NETs visualization and quantification

Polymorphonuclear cells (**PMN**; 2×10^5^) were attached to coverslips for 30 minutes at 37 °C. Cells were then stimulated with 5×10^−8^ M PMA and incubated for three hours at 37 °C. To visualize NETs, the cells were stained with mouse anti-dsDNA (Abcam, Cambridge, MA, USA), rabbit antihuman Neutrophil Elastase (Abcam, Cambridge, MA, USA), goat anti-rabbit Alexa Fluor 555, and goat anti-mouse Alexa Fluor 488 (Abcam, Cambridge, MA, USA). Samples were mounted using Fluoroshield mounting media (Abcam, Cambridge, MA, USA). The slide images were acquired using the Nikon Eclipse Ti (Zeiss LSM 800 confocal microscope) and analyzed with NIS-Elements (Nikon) or Fiji (NIH open source). For quantifying, the images were analyzed with Fiji, the number of cells ejecting extracellular DNA were counted, then the percentage of cells producing NETs were calculated per field of view (FOV), and then per each donor.

### Analysis of CFTR protein by immunofluorescence

For CFTR detection 10^5^ human neutrophils were seeded in coverslips coated with poly-L-lysine (Sigma Aldrich, St. Louis, MO). Cells were fixed, permeabilized, and stained with combined mouse anti-human CFTR clones CF3 or TJ-A9 (ThermoFischer Scientific), wheat germ agglutinin Alexa Fluor 647 or 555 (ThermoFischer Scientific), goat anti-mouse Alexa Fluor 488 (Abcam, Cambridge MA), rabbit anti-LAMP1, goat anti-rabbit IgG Alexa Fluor 647 and 4’,6-diamidino-2-phenylindole (**DAPI**) (ThermoFischer Scientific). Slides were imaged by confocal microscopy (Zeiss LSM 800 confocal microscope) and analyzed with NIS-Elements or Fiji.

### Flow cytometry

For analyzing CFTR by flow cytometry, non-CF or CF neutrophils were purified and stained in two different ways, non-permeabilized or permeabilized. The PMNs were stained with anti-human CFTR (clone TJA9) and anti-mouse IgG Alexa Fluor 488. Flow cytometry (LSR Fortessa, BD Biosciences) collected the cells, and FlowJo V10.10 and GraphPad Prism V10 were used for fluorescence and statistical analyses respectively.

### Electrophysiology recordings (patch-clamp in whole-cell configuration)

The extracellular solution was prepared with 145 mM NaCl, 15 mM sodium glutamate, 4.5 mM KCl, 1 mM MgCl_2_, 2 mM CaCl_2_, 10 mM HEPES, 5 mM glucose at pH 7.4. The intracellular solution was prepared at pH 7.2 with 139 mM CsCl, 2 mM MgCl_2_, 5 mM ethylene glycol tetraacetic acid, 10 mM HEPES, 5 mM glucose, 2 mM ATP, and 0.1 mM guanosine 5’-triphosphate. The CFTR current was induced in PMN using a cocktail of 15 μM forskolin, 100 μM IBMX, and 2 mM ATP. Some neutrophils were pretreated with Lumacaftor/Ivacaftor (5 μM / 5 μM) for one hour before the experiment at 37 °C. For inhibiting CFTR, 20 μM CFTR_inh-172_ or 10 μM PPQ-102 were added to the PMN. Patch-clamp experiments were conducted at room temperature using an Axopatch 200B and Multiclamp 700 B amplifiers (Molecular Devices Corp. CA, USA) equipped with pCLAMP 10.6 and 11 software and EPC 10 USB HEKA amplifier (HEKA Elektronik, Germany) and patch mater 10 (HEKA Elektronik, Germany). The recording electrode was made from borosilicate glass (World Precision Instruments, Inc, Sarasota FL, USA) with a Sutter puller (model P-97, Sutter Instruments, USA). The resistance was 3-5 mΩ. Three different protocols elicited whole cell currents: A) Voltage-step. From -100 mV to +100 mV, ΔV=10 mV, holding potential of -30 mV and 200 ms of duration. B) Voltage-step. From -100 mV to +100 mV, ΔV=20 mV, holding potential of -50 mV and 1000 ms of duration. C) Voltage-ramp protocol. From -100 mV to +100 mV, holding potential of 0 mV and 1000 ms of duration.

### Statistical analysis

Parametric unpaired Student’s *t*-test or nonparametric ANOVA, ONE-way test, statistically evaluated as bar graphs, according to each case. GraphPad Prism software (San Diego CA) performed all analyses. A value of *p*<0.05 was considered statistically significant.

## RESULTS

### CFTR is functionally expressed in neutrophils

Previously, we determined that neutrophils exhibiting F508del CFTR mutation were markedly impaired in antimicrobial killing, likely due to a reduced oxidative burst and a delayed NETosis [15]. Here, we first sought to confirm CFTR functional expression in neutrophils. To this end, we collected peripheral blood neutrophils from healthy donors (**non-CF**) and analyzed the mRNA expression of CFTR by RT-PCR. The mRNA was present in non-CF neutrophils (**Figure 1A)**. Next, immunoprecipitation analyzed for CFTR protein in non-CF. As expected, the glycosylated CFTR protein (β170 KDa) was detected in neutrophils (**Figure 1B**). Additionally, we performed an immunofluorescence to evaluate CFTR localization by using anti-CFTR (CF3), which binds to the first extracellular loop of the CFTR protein, wheat germ agglutinin to stain the plasma membrane, and DAPI for counterstaining the DNA. The expression of CFTR colocalized with the plasma membrane was observed in non-permeabilized neutrophils (**Figure 1C**). Total CFTR was visualized after non-CF neutrophils were permeabilized and stained. CFTR was colocalizing with the plasma membrane and Endoplasmic Reticulum (**ER**) (**Figure 1D**), indicating the presence of the CFTR protein in both the plasma membrane and the ER. Additionally, we analyzed CFTR expression in lysosomes (LAMP-1). We detected CFTR colocalized with both the plasma membrane and lysosomes (**Sup Fig 1A**).

**Figure 1.**
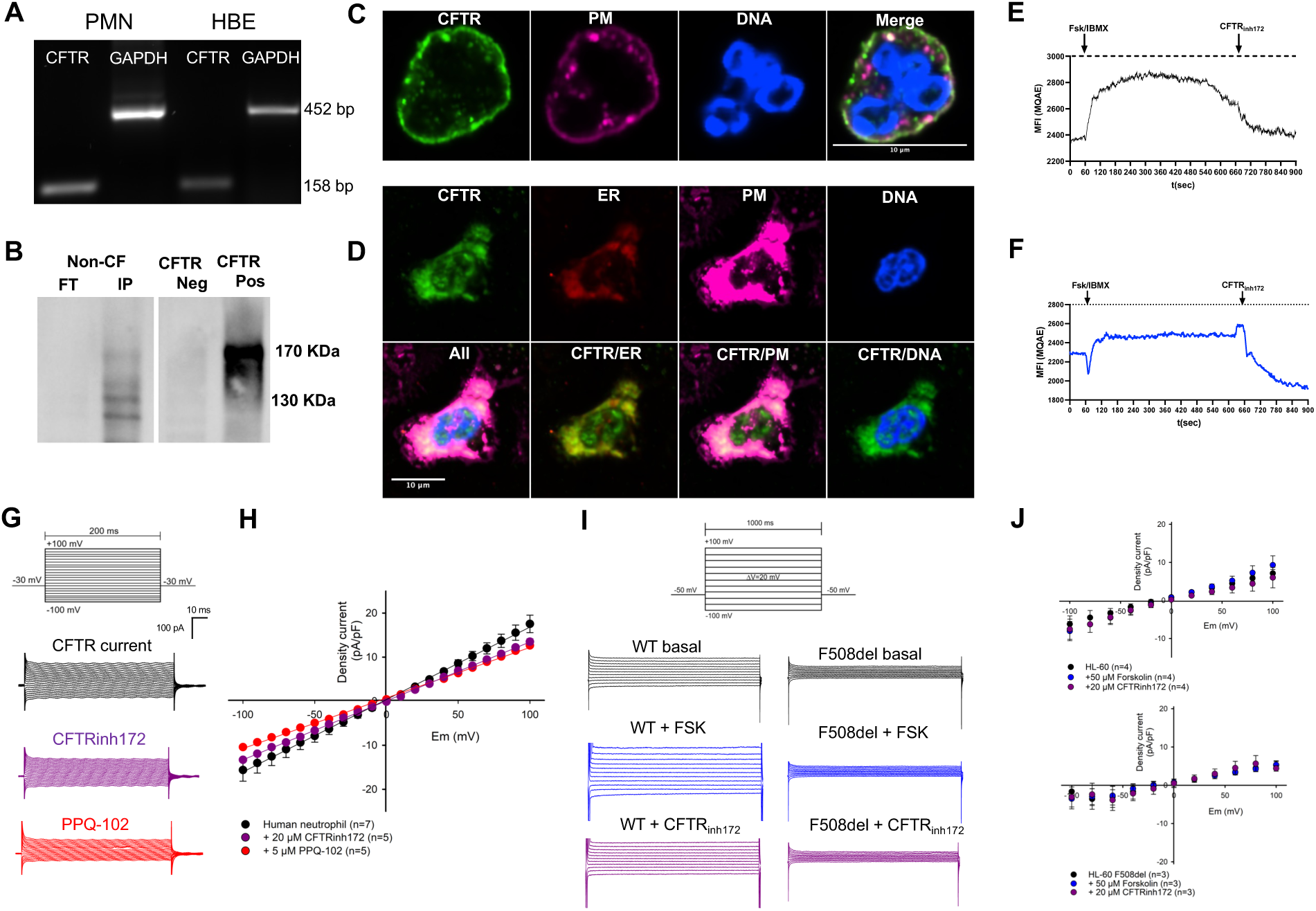
Functional expression of CFTR in neutrophils. **(A)** Non-CF neutrophils and human bronchial epithelial (HBE) cells were lysed, and RT-PCR assay for detecting Cystic Fibrosis Transmembrane-Conductance Regulator (CFTR) was performed. Glyceraldehyde-3-phosphate dehydrogenase (GAPDH) was run as a housekeeping gene. **(B)** Immunoprecipitation was run with non-CF neutrophil lysates; the immunoblot was stained with anti-CFTR (clone TJA9). **(C)** Non-CF blood neutrophils were fixed and stained with mouse anti-human CFTR clone CF3 (green). Wheat germ agglutinin (WGA) stained the plasma membrane, and DAPI stained DNA. **(D)** Some neutrophils were permeabilized and stained with anti-CFTR clone CF3 (green), WGA (magenta), anti-P4HB (endoplasmic reticulum, red). DAPI stained DNA. The colocalization between CFTR and ER, plasma membrane, and nucleus is displayed. The white bar represents 10 μm. **(E)** The kinetics of Cl^−^ efflux were acquired in neutrophils stained with MQAE. The intracellular Cl^−^ basal levels were recorded for 30 seconds, followed by stimulation with 20 μM forskolin/50 μM IBMX up to 660 seconds, then 20 μM CFTRinh-172 was added up to 900 seconds. **(F)** The halide assay was performed on neutrophils; the cells were stained with MQAE, followed by flow cytometry recording iodide. Non-CF neutrophils revealed halide efflux when CFTR was activated with forskolin/IBMX. **(G)** Representative CFTR current recordings by the patch-clamp in whole-cell configuration in non-CF neutrophils (black line), after adding 20 μM CFTRinh-172 (magenta) or 10 μM PPQ-102 (red). **(H)** I-V curves of CFTR currents after adding Forskolin/IBMX and PPQ-102 or CFTRinh-172. (5-7 cells, data expressed as mean ± s.e.m.). **(I)** CFTR currents of HL-60 cells were acquired by patch-clamp: basal levels (black), after adding forskolin (blue), and the CFTRinh-172 (magenta) in WT and F508del cells. **(J)** The I/V curves of the CFTR currents indicate a reduced response of F508del cells. All these measurements corroborated that CFTR is expressed and functional in neutrophils.

To analyze whether the CFTR is functional in neutrophils, we performed a CFTR-dependent Cl^−^ efflux assay. Neutrophils were loaded with the chloride-sensitive dye N-(ethoxycarbonylmethyl)-6-methoxyquinolinium bromide (MQAE), as we previously reported [28]. Kinetic recording of the basal intracellular Cl^−^ levels (**Figure 1E**) was followed by adding forskolin/IBMX, which activates adenylate cyclase, producing cAMP [30–32] to induce CFTR activation, and finally adding the CFTR_inh-172_ to block CFTR-dependent Cl^−^ efflux. Since CFTR can mobilize I^−^, the Cl^−^ was depleted in the media and replaced by I^−^ ions. The standard halide assay modified by flow cytometry [28], measured the iodide efflux (**Figure 1F**). The kinetics revealed iodide efflux once forskolin/IBMX was added to the neutrophils. Adding CRTR_inh-172_ to the media quenched the fluorescence.

Next, we performed whole-cell patch clamp to measure the CFTR current in neutrophils. We recorded the baseline CFTR-dependent current in non-CF neutrophils (**Figure 1G**). Moreover, adding the CFTR_inh-172_ or PPQ-102 drastically reduced the CFTR-dependent current (23.04% with CFTR_inh-172_, 27.91% with PPQ-102). The IV plot illustrates the traces of the CFTR current basally and with CFTR inhibition (**Figure 1H),** the reversal potential shifts to 0 mV. However, the ionic recording conditions are not symmetrical; that effect could contribute Na^+^ flux through sodium channels like ENaC as previously reported [33].

Further, we used the neutrophil-like HL-60 F508del cells [34], as a CFTR-deficient cell model to analyze CFTR-dependent mechanisms. We analyzed the glycosylated CFTR protein expression in undifferentiated cells and differentiated cells treated with DMSO or retinoic acid for 5 days before use [34]. The undifferentiated F508del cells exhibited a reduced matured CFTR band (≈170 kDa) as compared to the WT cells (**Sup Fig 1B**). The F508del DMSO or retinoic acid differentiated cells did not display a mature CFTR band as compared to the WT cells. Additionally, we analyzed whether the CFTR current could be detected in CFTR-deficient cells. The CFTR current was present in basal conditions and after the stimulation with forskolin in WT cells (**Figure 1I, J**); however, the CFTR current was deeply reduced in F508del mutant cells. No effect was observed after adding the CFTR_inh-172_ to the F508del cells. Together, these results confirm that the CFTR is present and functional in healthy human neutrophils, but CFTR function is not present in F508del mutant neutrophils.

### ETI treatment restores the expression and function of the CFTR in CF neutrophils

To analyze the effect of ETI on the expression and function of CFTR in neutrophils, we used WT or F508del HL-60 differentiated cells. The cells were treated with ETI and stained with anti-CFTR (non-permeabilized). The F508del cells did not indicate CFTR expression in the plasma membrane (**Figure 2A**). However, treating with ETI restored CFTR expression in F508del cells, and similar results were observed when using undifferentiated HL-60 cells (**Sup Fig 2A**).

**Figure 2.**
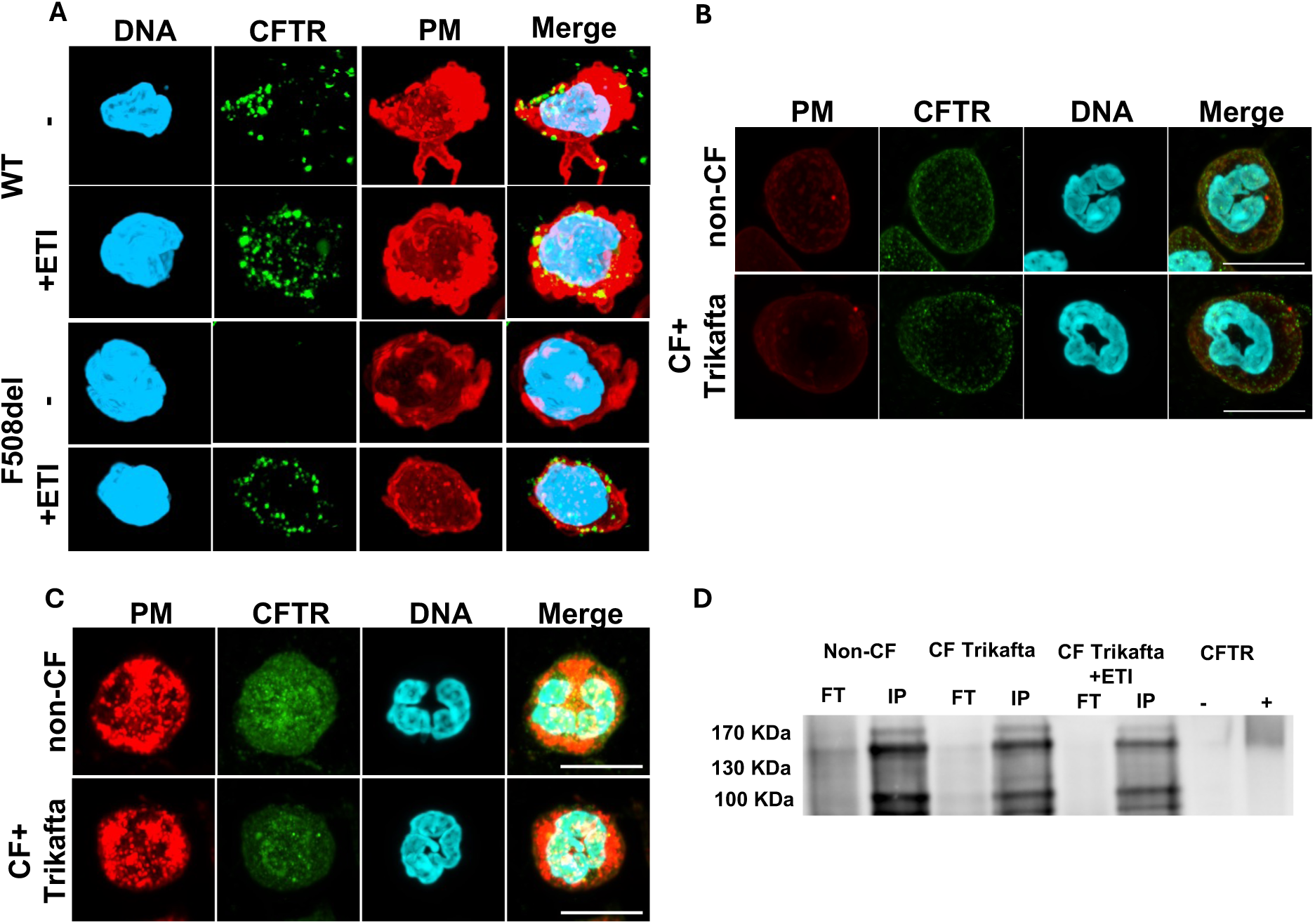
ETI treatment restores CFTR localization in CF neutrophils. **(A)** WT and F508del HL-60 cells differentiated with DMSO to neutrophil-like cells, were treated with 5 µM Elexacaftor/5 µM Tezacaftor/1 µM Ivacaftor (**ETI**) and stained with anti-CFTR (green), WGA (plasma membrane, red) and DAPI (blue), F508del cells treated with ETI recovered CFTR (n=3) expression. Non-permeabilized **(B)** or permeabilized **(C)** neutrophils from pwCF under treatment with Trikafta^®^ (ETI) were stained with anti-CFTR (green), and counterstained with DAPI (blue) and WGA (red). **(D)** Immunoprecipitation was run with non-CF, CF neutrophil lysates; some cells were pretreated with ETI. The immunoblot was stained with anti-CFTR (clone TJA9), and the CFTR mature band displays at β170 KDa. Treating with ETI in vivo and in vitro modified the cellular location of the CFTR in non-CF and CF neutrophils.

We also analyzed the effect of ETI on CFTR expression in non-CF or CF neutrophils. The cells were exposed to ETI in vitro and then stained with anti-CFTR. ETI treatment slightly increased CFTR expression in the plasma membrane of non-CF and CF neutrophils (**Sup Fig 2B-C**). Furthermore, neutrophils from pwCF under treatment with ETI were collected and stained with anti-CFTR. ETI-treated CF neutrophils revealed slightly lower CFTR expression in the plasma membrane than non-CF neutrophils in non-permeabilized cells (**Figure 2B**). At the same time, the total CFTR was similar in non-CF and CF neutrophils in permeabilized cells (**Figure 2C**). Additionally, we performed flow cytometry to quantify CFTR expression and found that both non-CF and CF neutrophils from pwCF under ETI treatment revealed similar mean fluorescent intensity corresponding to CFTR in the plasma membrane (**Sup Fig 2D**).

We also analyzed the glycosylated CFTR protein expression in CF neutrophils treated with ETI (**Figure 2D**). We detected slightly lower levels of the CFTR matured protein in CF neutrophils as compared with non-CF, and similar observations were seen in CF neutrophils re-treated with ETI in vitro as well.

Next, we collected peripheral blood neutrophils and performed patch clamp in whole-cell configuration to measure CFTR Cl^−^ currents. CF neutrophils collected from pwCF treated with ETI exhibited a reduced Cl^−^ current as compared to non-CF neutrophils (**Figure 3A**). To confirm the effect of CFTR modulators in vitro on the CFTR-dependent current, we treated non-CF neutrophils with the modulator combination Lumacaftor/Ivacaftor (LI), followed by measuring the Cl^−^ current (**Figure 3B**). The CFTR modulators drastically increased the density current in neutrophils (∼20 pA/pF to ∼150 pA/pF, at +100 mV).

**Figure 3.**
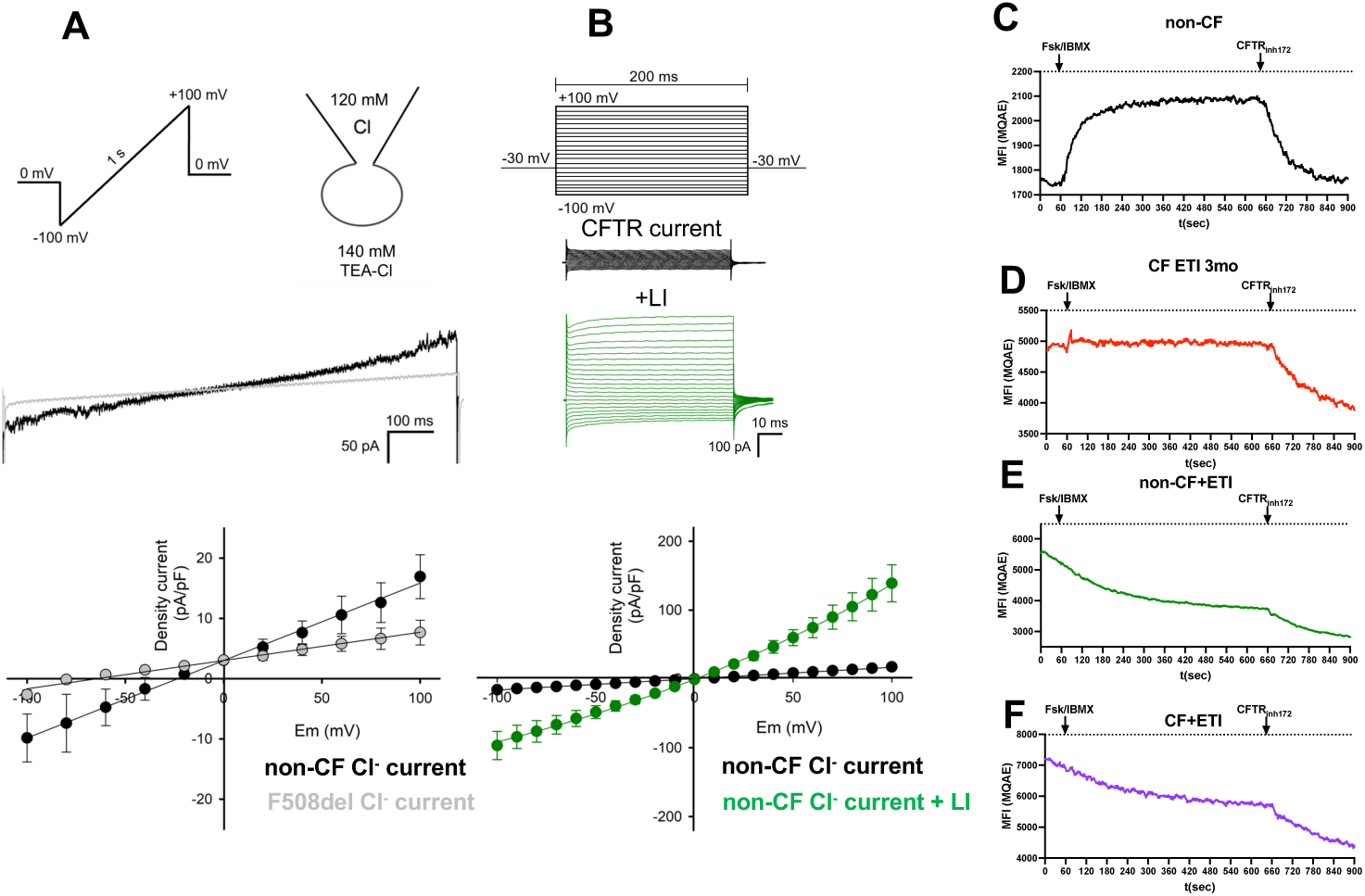
CFTR modulators restore the CFTR function in CF neutrophils. Chloride currents were recorded in CF neutrophils by patch-clamp in whole-cell configuration **(A)**. The CFTR current was analyzed in CF neutrophils from pwCF under treatment with Trikafta^®^ (grey line) or non-CF neutrophils (black line). CF neutrophils showed a reduced Cl^−^ current as compared to non-CF neutrophils. **(B)** Representative voltage-step recordings and I-V curves of CFTR current in non-CF neutrophils (black line) and after adding 5 μM Ivacaftor/Lumacaftor (green line) for 1 h as determined in 3-8 cells (Data expressed as mean ± s.e.m.). Chloride efflux kinetics recorded by flow cytometry were performed in non-CF neutrophils **(C)** or CF neutrophils from patients under treatment with Trikafta^®^ for 3 mo **(D). (E**) Non-CF neutrophils or **(F)** CF neutrophils (under Trikafta^®^) were treated for 1 h with ETI, followed by the Cl^−^ efflux measurements. Neutrophils treated with CFTR modulators exhibited restored CFTR function.

Furthermore, the effect of ETI on the Cl^−^ efflux in neutrophils was analyzed. Flow cytometric measurements of Cl^−^ efflux in non-CF neutrophils displayed significant CFTR-dependent Cl^−^ efflux activation (**Figure 3B**). Notably, CF neutrophils from patients treated with ETI revealed an increased fluorescence at baseline, indicating a lower intracellular Cl^−^ concentration than non-CF neutrophils (**Figure 3D**). Additionally, CF neutrophils did not further respond to stimulation with forskolin/IBMX but exhibited a substantial reduction in fluorescence once the CFTR_inh-172_ was added to the cells. It is possible that continuous exposure of CF neutrophils to modulators in vivo increased CFTR mobilization to the plasma membrane and constant CFTR mediated Cl^−^ efflux. Similarly, non-CF neutrophils or CF neutrophils treated *in vitro* with ETI for one hour displayed increased levels of fluorescence at baseline, which correlates with Cl^−^ efflux (**Figure 3E, F).** Moreover, neither non-CF nor CF neutrophils pretreated with ETI further responded to stimulation with forskolin/IBMX, but adding the CFTR inhibitor reduced the fluorescence, indicating replenished intracellular Cl^−^ accumulation.

### ETI treatment restores the defective antimicrobial mechanisms of CF neutrophils

CF neutrophils were treated with single modulators Ivacaftor, Lumacaftor, or Tezacaftor for one hour before infection with *B. cenocepacia,* and intracellular killing was evaluated three hours postinfection. Treatment with single CFTR modulators modestly increased the intracellular antimicrobial killing of CF neutrophils (**Figure 4A**). In contrast, the pretreatment with combinations of Tezacaftor/Ivacaftor (TI) or Elexacaftor/Tezacaftor/Ivacaftor (**ETI**), markedly potentiated the antimicrobial killing of CF neutrophils (**Figure 4B**). Ex vivo neutrophils isolated from pwCF under treatment for 3 months with Symdeko® (TI) or Trikafta^®^ (ETI) were collected and challenged with *B. cenocepacia* (**Figure 4C, D**). CF neutrophils from patients under Symdeko® did not exhibit significant recovery in their intracellular antimicrobial killing. Interestingly, CF neutrophils from patients under treatment with Trikafta^®^ fully recovered their antimicrobial response.

**Figure 4.**
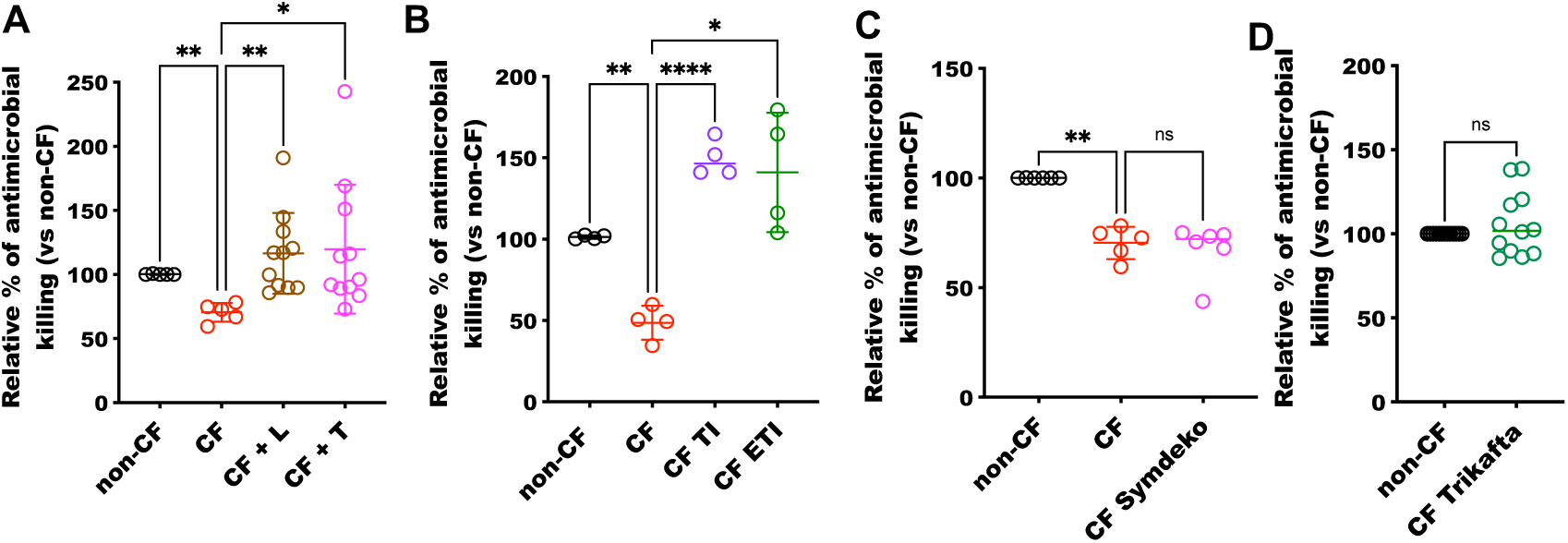
ETI treatment restores the antimicrobial response in CF neutrophils. Non-CF and CF neutrophils were treated with **(A)** Lumacaftor, Tezacaftor, or **(B)** Ivacaftor/Tezacaftor or Ivacaftor/Tezacaftor/Elexacaftor for 1 h and infected with *B. cenocepacia* (MOI = 10). The intracellular antimicrobial killing was analyzed after 3 h. Brown-Forsythe and Welch ANOVA test multiple comparison were applied for the statistical analysis (n=5, *=p<0.05, **=p<0.01, ***=p<0.001). Non-CF and CF neutrophils from patients with or without treatment with **(C)** Symdeko® (Ivacaftor/Tezacaftor, TI) or **(D)** Trikafta^®^ (Elexacaftor/Tezacaftor/Ivacaftor, ETI) for 3 mo were infected with *B. cenocepacia* (MOI = 10), and intracellular antimicrobial killing was analyzed after 3 h. Brown-Forsythe and Welch ANOVA test multiple comparison or Welch’s t test were applied for the statistical analysis (n=5-12, **=p<0.01, ns=p>0.05).

Since treating with ETI restored the intracellular antimicrobial killing of CF neutrophils, we analyzed the potential pathways responsible for this recovery. We focused first on the NADPH oxidase pathway. To this end, we measured ROS production in non-CF neutrophils and CF neutrophils by stimulating the protein kinase C pathway with PMA. CF neutrophils produced less ROS than non-CF neutrophils (**Figure 5A**). CF neutrophils from a pwCF treated with ETI, displayed increased ROS production, as compared to the CF neutrophils without treatment (**Figure 5B**). The ROS kinetics in CF neutrophils treated with ETI were very similar to those of non-CF neutrophils. The quantitative analysis compared the area under the curve of the ROS kinetics (**Sup Fig 3A-B**). We also measured hypochlorite production in non-CF and CF neutrophils from pwCF under ETI treatment, CF neutrophils with Trikafta^®^ treatment displayed greater levels of hypochlorite than non-CF neutrophils. However, when non-CF or CF neutrophils were treated in vitro with ETI, hypochlorite production was reduced in both cases (**Sup Fig 3C**). Additionally, we measured the oxidative stress produced by oxidizing DNA in neutrophils, non-CF and CF neutrophils from pwCF, under ETI treatment, which exhibited similar levels of oxidative stress (**Figure 5C**).

**Figure 5.**
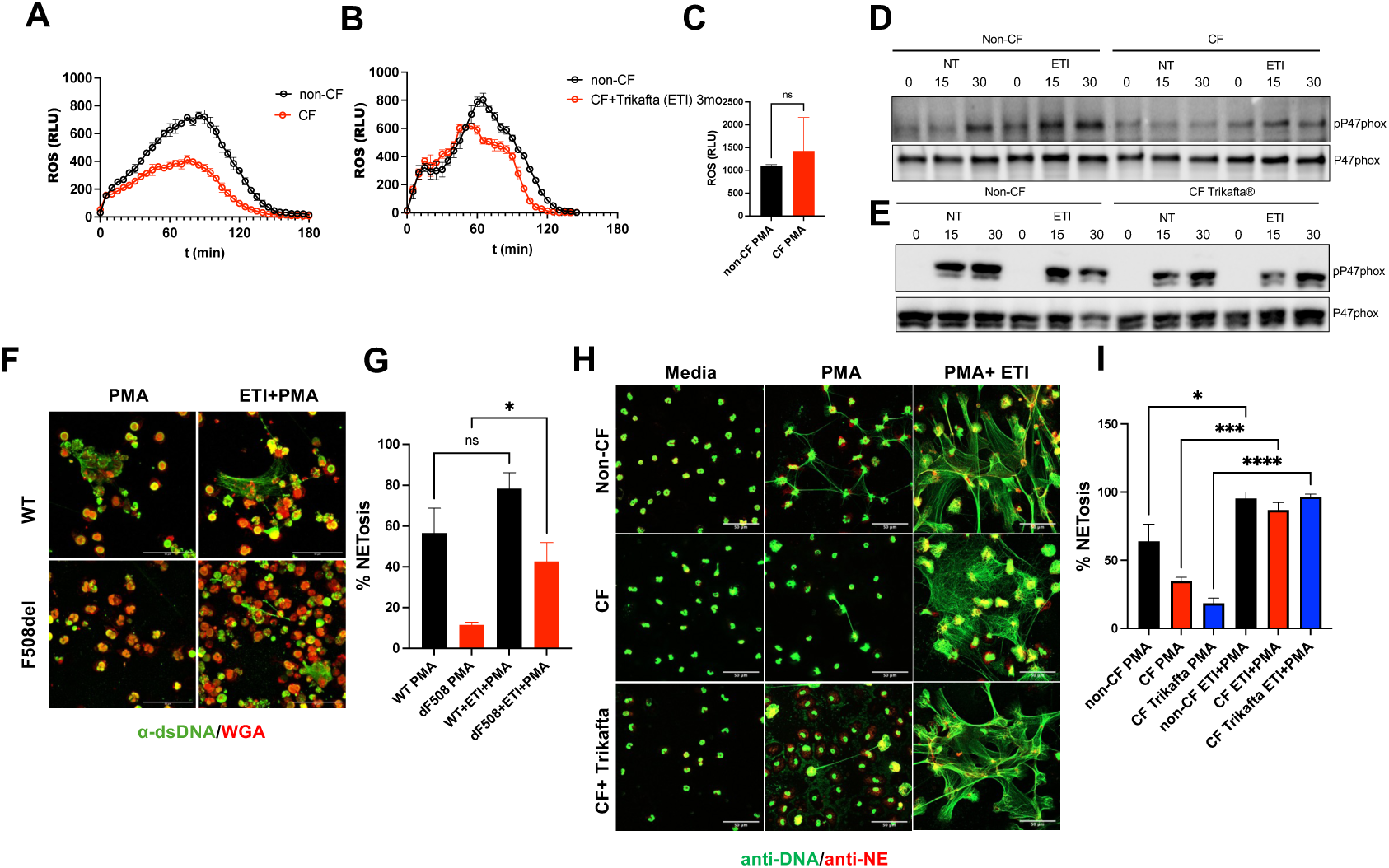
ETI promotes the oxidative burst and NETosis in CF neutrophils. **(A)** The oxidative response was measured in non-CF and CF neutrophils as a kinetic rate recorded after adding 50 nM PMA. The oxidative response was detected with luminol, and the area under the curve was quantified, (****=p>0.001, Welch’s t-test). **(B)** CF neutrophils treated with Trikafta® improved the oxidative response. **(C)** DNA oxidation was measured in non-CF and CF neutrophils from pwCF under Trikafta^®^ treatment. The cells were stimulated for 30 minutes with PMA, and the fluorescence was expressed as Relative Light Units (RLU) (ns=p<0.05, Welch’s t test). For analyzing the NADPH oxidase complex, **(D)** non-CF or CF neutrophils were stimulated with 50 nM PMA for 15 or 30 minutes; some cells were pretreated with ETI before the stimulation. After cells were lysed, western blot measured phosphorylated P47^phox^ expression or total P47^phox^. **(E)** P47^phox^ phosphorylated or total P47^phox^ were measured on CF neutrophils under treatment with Trikafta^®^. CF neutrophils partially restore their activation levels of Nox2 in pwCF under treatment with Trikafta^®^. **(F)** HL-60 cells differentiated with DMSO to neutrophil-like cells were pretreated with ETI and stimulated with PMA for 3 h, the NETs were stained with anti-dsDNA (green) and WGA (red), and **(G)** the percentage of cells producing NETs were quantitated. **(H)** Non-CF and CF neutrophils were seeded and treated with ETI, then cells were stimulated with PMA and incubated for 3 h. After fixation and permeabilization, the cells were stained with mouse anti-dsDNA (green), anti-Neutrophil Elastase (red), and the images were acquired by confocal microscopy and are representative of three independent experiments, **(I)** the NETs were quantitated and analyzed (*=p<0.5, ***=p<0.01, ****=p<0.001, one way ANOVA)

To further evaluate NADPH oxidase activation, we determined phosphorylation of the subunit phos-p47^phox^ in CF and non-CF neutrophils. Non-CF neutrophils exhibited higher levels of phosp47^phox^ than CF neutrophils. Pretreatment with ETI clearly augmented the levels of phos-p47^phox^ in both non-CF and CF neutrophils (**Figure 5D**). Finally, CF neutrophils collected from pwCF under treatment with ETI reveal lower levels of phos-p47^phox^ than non-CF neutrophils (**Figure 5E**).

### CFTR modulators promote NETosis in CF neutrophils

The ultimate resource of neutrophils in their attempts to kill bacteria is developing cell death through NETosis. This cellular process is mainly driven by activating the NADPH oxidase pathway [35]. We previously established that CF neutrophils exhibit a delayed NETosis process [15]. Thus, we reasoned that if ETI restores CFTR signaling in CF neutrophils, then ETI treatment could reestablish NETosis in CF neutrophils.

We found that differentiated HL-60 WT neutrophil-like cells stimulated with PMA for three hours could produce NETs. However, HL-60 F508del neutrophil-like cells exhibited fewer structures of NETs (**Figure 5F**). Treating F508del in vitro with ETI restored NET production. Semi-quantifying NETs established a correction in NET production in the F508del cells (**Figure 5G**). Additionally, non-CF neutrophils stimulated with PMA for three hours ejected DNA decorated with the granular protein Neutrophil Elastase (NE) and ETI potentiated NET-release in CF and non-CF neutrophils (**Figure 5H).** Like our previous findings, CF neutrophils did not reveal the same NETs production as non-CF neutrophils; however, in vitro treatment with CFTR modulators completely re-established or even increased the process of NETosis. Semi-quantifying NETs established a pronounced NETs production in CF neutrophils, which was not observed in non-CF and CF neutrophils from pwCF under ETI treatment (ex vivo) (**Figure 5I**).

## DISCUSSION

Chronic lung inflammation has been a significant concern for pwCF, leading to bronchiectasis and a decline in lung function over time. One major topic of interest in the CF field is whether lung inflammation results from changes in aberrant epithelial cell function and mucus secretion, intrinsic defects in CF immune cells, or a combination thereof. Extrinsic lung environmental factors include accumulating a thick and dense mucus that leads to airway obstruction and reduced oxygen levels, causing hypoxia and sterile inflammation [36]. Additionally, the thick layer of mucus promotes biofilms by pathogens associated with pwCF, including *Pseudomonas aeruginosa* [37], *Burkholderia cenocepacia,* and *Staphylococcus aureus* [11]. In contrast, intrinsic factors are directly associated with CFTR dysfunction in the immune cells by promoting hyperinflammation due to excessively producing inflammatory cytokines [38], and a diminished antimicrobial response [15, 24]. While CFTR expression has been associated with neutrophils, some reports suggest that CFTR is mainly stored in phagocytic vesicles [39–41]. In this study, we wanted to unequivocally confirm the CFTR protein expression in the plasma membrane and its impact on the basic physiology and antimicrobial response of neutrophils. We indeed established CFTR protein expression in neutrophils at mRNA and protein levels, as the CFTR protein was detected in ER compartments and the plasma membrane. Additionally, we demonstrated by patch clamp electrophysiology CFTR channel activity in the plasma membrane of neutrophils for the first time. The CFTR-mediated current was detected in non-CF neutrophils, but it was absent in CFTR mutant cells. We also corroborated that CFTR is functional in neutrophils by using the halide assay method modified for cells in suspension and recorded by flow cytometry [28].

Previous reports have associated deficient CFTR function with reduced reactive oxygen species (ROS) production [15], and dysregulated hypochlorous acid (HClO^−^) production [15, 42]. CFTR-mediated transport of Cl^−^ ions has been proposed to contribute to HClO^−^ formation in phagolysosomes through the myeloperoxidase pathway in healthy neutrophils [43–45].

During the past few years, most of the efforts to develop CFTR modulator therapies have focused on correcting CFTR transport to the plasma membrane in pwCF carrying F508del mutation and increasing the flow of ions through activated CFTR present at the cell surface in pwCF carrying dysfunctional CFTR. The changes induced by ETI at the cellular level have been analyzed mainly in airway epithelial cells [21–23]. In contrast, few studies have investigated whether ETI restores functional CFTR expression in phagocytic cells. This lack of research is in part due to inconclusive reports regarding CFTR expression at the plasma membrane of these cells. Addressing this knowledge gap, we previously reported, in collaboration with Dr. Kopp’s group, that treating CF macrophages with ETI markedly increased the functional expression of CFTR in the plasma membrane. CFTR correction in CF macrophages correlated with improved phagocytosis and better clinical outcomes [24]. However, the effect of ETI on neutrophils has not been investigated.

The improvement in intracellular antimicrobial killing in CF neutrophils treated with CFTR modulators observed in this work could be explained, at least in part, by the fact that modulator treatments increased the density of the CFTR protein in the plasma membrane and intracellular compartments, including the phagosomes. This increment in membranal CFTR could restore the Cl^−^ flux to the intracellular compartments to produce HClO^−^, which is one of the most effective antimicrobial agents in neutrophils.

Another important antimicrobial mechanism in CF neutrophils that is significantly improved upon treating with ETI is NETs production. Two types of NETosis are referred as suicidal NETosis, which is NADPH oxidase-dependent, and vital NETosis, which is NADPH oxidase-independent [46]. In the suicidal NETosis, once the NADPH oxidase is activated, ROS are released, including O_2_, H_2_O_2_, and HClO^−^ [47]. Contributing NETosis to the aggravated pathology in the lung of pwCF is controversial as some authors claim that CF neutrophils are more prone to develop excessive NETs [48]. In contrast, our previous data revealed that CF neutrophils developed delayed and inefficient NETosis when the NADPH oxidase pathway is induced [15]. Moreover, here we demonstrate that treating CF neutrophils with CFTR modulators fully restores effective NETs production when inducing NADPH oxidase pathway. The potential mechanism that restores NETosis in CF neutrophils after treatment with CFTR modulators may be attributed to the increased phosphorylation of P47^phox^. This enhancement likely reinstates NOX2 activity and boosts the NADPH oxidase final products, including HClO^−^. Recovering NADPH oxidase function and suicidal NETosis may transform the CF neutrophils into a more efficient antimicrobial barrier, which will likely correlate with resolving infection in pwCF better, along with reducing tissue damage in the lung. This idea is consistent with the fact that treating pwCF with Trikafta^®^ reduces the frequency and severity of lung infections, improves FEV_1_, reduces the sweat Cl^−^ concentration, and enhances the quality of life overall in pwCF [20].

In summary, here we analyzed the effects of ETI in CF neutrophils. We observed an increased CFTR protein expression in the plasma membrane of CF neutrophils, which correlated with CFTR-mediated current after CFTR modulator treatment. ETI treatment also resulted in reducing the intracellular Cl^−^ levels in CF neutrophils and improving the intracellular antimicrobial killing by restoring the NADPH oxidase activity and Neutrophil Extracellular Traps (NETs) production.

Together, the data presented strongly suggest that CF neutrophils’ dysfunction is an intrinsic defect related to CFTR mutation. However, once CFTR modulators correct CFTR expression and function, increasing the density of the mature CFTR ion channel in the plasma membrane fixes the defective antimicrobial mechanism in CF neutrophils, which thus restores the NADPH oxidase pathway. In turn, this pathway potentiates the intracellular antimicrobial killing and produces NETs.

In conclusion, restoring Cl^−^ ion imbalances in CF neutrophils appears to be essential for effective antimicrobial functions in these cells and offers the possibility of testing new pharmacologic molecules and alternative treatments targeting ion channels in pwCF. Re-establishing ion homeostasis in CF neutrophils of pwCF that are not candidates to receive the HEMT (type I mutations) could improve their life expectancy and quality of life.

### Limitations of the study

In this work, we focused on studying how the CFTR-deficiency impacts in the antimicrobial response of neutrophils. We analyzed samples from pwCF carrying F508del mutation (type II mutation), which are eligible to take CFTR modulators, and we did not study pwCF with other CFTR mutations. Our experiments were performed by using peripheral blood neutrophils collected from healthy donors (non-CF) or pwCF. Although our overall research goal was to elucidate intrinsic dysfunction in blood CF neutrophils, regretfully we had no access to neutrophils from sputum or bronchoalveolar lavages for possible comparisons. Additionally, we only analyzed the antimicrobial mechanism against *Burkholderia cenocepacia*, but no against other CF pathogens.

Nonetheless, our data show strong evidence of the importance of the CFTR ion channel on promoting an efficient antimicrobial response mediated by intracellular and extracellular mechanisms.

## Acknowledgments

Human blood samples were provided through the Cure CF Columbus Translational and Data Core (C3TDC). C3TDC is supported by the Division of Pediatric Pulmonary Medicine, the Biopathology Center Core, and the Data Collaboration Team at Nationwide Children’s Hospital. Grant support provided by The Ohio State University Center for Clinical and Translational Science (National Center for Advancing Translational Sciences, Grant UL1TR002733) and by the Cystic Fibrosis Foundation (Research Development Program, Grant MCCOY24RO). We acknowledge Dr. Melody Davis for critically reading and editing the manuscript, Stephanie Sliemers from the C3TDC for providing patient’s demographic information and the microscopy core at Nationwide Children’s Hospital.

## Authors’ contributions

F.R.A and S.P.S. conceptualized the study and designed the experiments. F.R.A., R.R., A.M.B. and V.L.C. performed the experiments. F.R.A., R.R., A.M.B. and V.L.C. analyzed data. F.R.A. wrote the manuscript draft. F.R.A., R.R., A.M.B., V.L.C., H.S., K.S.M, B.T.K. and S.P.S. revised the manuscript. S.P.S. supervised the study. All authors have approved the submitted version of the study and their contributions.

## REFERENCES

1. Shteinberg, M., et al., Cystic fibrosis. Lancet, 2021. 397(10290): p. 2195–2211.

2. Rommens, J.M., et al., Identification of the cystic fibrosis gene: chromosome walking and jumping. Science, 1989. 245(4922): p. 1059–65.

3. Riordan, J.R., et al., Identification of the cystic fibrosis gene: cloning and characterization of complementary DNA. Science, 1989. 245(4922): p. 1066–73.

4. Kerem, B., et al., Identification of the cystic fibrosis gene: genetic analysis. Science, 1989. 245(4922): p. 1073–80.

5. Bacalhau, M., et al., Elexacaftor-Tezacaftor-Ivacaftor: A Life-Changing Triple Combination of CFTR Modulator Drugs for Cystic Fibrosis. Pharmaceuticals (Basel), 2023. 16(3).

6. Jia, S. and J.L. Taylor-Cousar, Cystic Fibrosis Modulator Therapies. Annu Rev Med, 2023. 74: p. 413–426.

7. Liu, F., et al., Molecular Structure of the Human CFTR Ion Channel. Cell, 2017. 169(1): p. 85–95 e8.

8. http://www.genet.sickkids.on.ca/StatisticsPage.html.

9. Lukacs, G.L. and A.S. Verkman, CFTR: folding, misfolding and correcting the DeltaF508 conformational defect. Trends Mol Med, 2012. 18(2): p. 81–91.

10. Elborn, J.S., Identification and management of unusual pathogens in cystic fibrosis. J R Soc Med, 2008. 101 **Suppl 1**(Suppl 1): p. S2–5.

11. Blanchard, A.C. and V.J. Waters, Opportunistic Pathogens in Cystic Fibrosis: Epidemiology and Pathogenesis of Lung Infection. J Pediatric Infect Dis Soc, 2022. 11(Supplement_2): p. S3–S12.

12. Lipuma, J.J., The changing microbial epidemiology in cystic fibrosis. Clin Microbiol Rev, 2010. 23(2): p. 299–323.

13. Nichols, D.P. and J.F. Chmiel, Inflammation and its genesis in cystic fibrosis. Pediatr Pulmonol, 2015. 50 **Suppl 40**: p. S39–56.

14. Pohl, K., et al., A neutrophil intrinsic impairment affecting Rab27a and degranulation in cystic fibrosis is corrected by CFTR potentiator therapy. Blood, 2014. 124(7): p. 999–1009.

15. Robledo-Avila, F.H., et al., Dysregulated Calcium Homeostasis in Cystic Fibrosis Neutrophils Leads to Deficient Antimicrobial Responses. J Immunol, 2018. 201(7): p. 2016–2027.

16. Cuevas-Ocana, S., et al., The era of CFTR modulators: improvements made and remaining challenges. Breathe (Sheff), 2020. 16(2): p. 200016.

17. Lopes-Pacheco, M., CFTR Modulators: The Changing Face of Cystic Fibrosis in the Era of Precision Medicine. Front Pharmacol, 2019. 10: p. 1662.

18. McCoy, K.S., et al., Clinical change 2 years from start of elexacaftor-tezacaftor-ivacaftor in severe cystic fibrosis. Pediatr Pulmonol, 2023. 58(4): p. 1178–1184.

19. Zaher, A., et al., A Review of Trikafta: Triple Cystic Fibrosis Transmembrane Conductance Regulator (CFTR) Modulator Therapy. Cureus, 2021. 13(7): p. e16144.

20. Barry, P.J., et al., Triple Therapy for Cystic Fibrosis Phe508del-Gating and -Residual Function Genotypes. N Engl J Med, 2021. 385(9): p. 815–825.

21. Laselva, O., et al., Rescue of multiple class II CFTR mutations by elexacaftor+tezacaftor+ivacaftor mediated in part by the dual activities of elexacaftor as both corrector and potentiator. Eur Respir J, 2021. 57(6).

22. Laselva, O. and M. Conese, Elexacaftor/Tezacaftor/Ivacaftor Accelerates Wound Repair in Cystic Fibrosis Airway Epithelium. J Pers Med, 2022. 12(10).

23. Veit, G., et al., Allosteric folding correction of F508del and rare CFTR mutants by elexacaftor-tezacaftor-ivacaftor (Trikafta) combination. JCI Insight, 2020. 5(18).

24. Zhang, S., et al., Cystic fibrosis macrophage function and clinical outcomes after elexacaftor/tezacaftor/ivacaftor. Eur Respir J, 2023. 61(4).

25. Cantin, A.M., et al., Inflammation in cystic fibrosis lung disease: Pathogenesis and therapy. J Cyst Fibros, 2015. 14(4): p. 419–30.

26. Casey, M., et al., Effect of elexacaftor/tezacaftor/ivacaftor on airway and systemic inflammation in cystic fibrosis. Thorax, 2023. 78(8): p. 835–839.

27. Schmidt, H., et al., Multimodal analysis of granulocytes, monocytes, and platelets in patients with cystic fibrosis before and after Elexacaftor-Tezacaftor-Ivacaftor treatment. Front Immunol, 2023. 14: p. 1180282.

28. Montanez-Barragan, A., et al., Flow cytometric measurement of CFTR-mediated chloride transport in human neutrophils. J Leukoc Biol, 2025.

29. Zhang, S., et al., Consequences of CRISPR-Cas9-Mediated CFTR Knockout in Human Macrophages. Front Immunol, 2020. 11: p. 1871.

30. Awad, J.A., et al., Interactions of forskolin and adenylate cyclase. Effects on substrate kinetics and protection against inactivation by heat and N-ethylmaleimide. J Biol Chem, 1983. 258(5): p. 2960–5.

31. Robbins, J.D., et al., Forskolin carbamates: binding and activation studies with type I adenylyl cyclase. J Med Chem, 1996. 39(14): p. 2745–52.

32. Parsons, W.J., V. Ramkumar, and G.L. Stiles, Isobutylmethylxanthine stimulates adenylate cyclase by blocking the inhibitory regulatory protein, Gi. Mol Pharmacol, 1988. 34(1): p. 37–41.

33. Reus-Chavarria, E., et al., Enhanced expression of the Epithelial Sodium Channel in neutrophils from hypertensive patients. Biochim Biophys Acta Biomembr, 2019. 1861(2): p. 387–402.

34. Wang, D., et al., ERK is involved in the differentiation and function of dimethyl sulfoxide-induced HL-60 neutrophil-like cells, which mimic inflammatory neutrophils. Int Immunopharmacol, 2020. 84: p. 106510.

35. Azzouz, D., M.A. Khan, and N. Palaniyar, ROS induces NETosis by oxidizing DNA and initiating DNA repair. Cell Death Discov, 2021. 7(1): p. 113.

36. Montgomery, S.T., et al., Hypoxia and sterile inflammation in cystic fibrosis airways: mechanisms and potential therapies. Eur Respir J, 2017. 49(1).

37. Moreau-Marquis, S., B.A. Stanton, and G.A. O’Toole, Pseudomonas aeruginosa biofilm formation in the cystic fibrosis airway. Pulm Pharmacol Ther, 2008. 21(4): p. 595–9.

38. Harwood, K.H., et al., Anti-Inflammatory Influences of Cystic Fibrosis Transmembrane Conductance Regulator Drugs on Lung Inflammation in Cystic Fibrosis. Int J Mol Sci, 2021. 22(14).

39. Painter, R.G., et al., CFTR Expression in human neutrophils and the phagolysosomal chlorination defect in cystic fibrosis. Biochemistry, 2006. 45(34): p. 10260–9.

40. Zhou, Y., et al., Cystic fibrosis transmembrane conductance regulator recruitment to phagosomes in neutrophils. J Innate Immun, 2013. 5(3): p. 219–30.

41. Wang, G. and W.M. Nauseef, Neutrophil dysfunction in the pathogenesis of cystic fibrosis. Blood, 2022. 139(17): p. 2622–2631.

42. Wang, G., Chloride flux in phagocytes. Immunol Rev, 2016. 273(1): p. 219–31.

43. Painter, R.G., et al., CFTR-mediated halide transport in phagosomes of human neutrophils. J Leukoc Biol, 2010. 87(5): p. 933–42.

44. Aiken, M.L., et al., Chloride transport in functionally active phagosomes isolated from Human neutrophils. Free Radic Biol Med, 2012. 53(12): p. 2308–17.

45. Ng, H.P., et al., Neutrophil-mediated phagocytic host defense defect in myeloid Cftr-inactivated mice. PLoS One, 2014. 9(9): p. e106813.

46. Huang, J., et al., Molecular mechanisms and therapeutic target of NETosis in diseases. MedComm (2020), 2022. 3(3): p. e162.

47. de Bont, C.M., et al., Stimulus-dependent chromatin dynamics, citrullination, calcium signalling and ROS production during NET formation. Biochim Biophys Acta Mol Cell Res, 2018. 1865(11 Pt A): p. 1621–1629.

48. Law, S.M., et al., Neutrophil extracellular traps are associated with airways inflammation and increased severity of lung disease in cystic fibrosis. ERJ Open Res, 2024. 10(6).

