## Supplemental Figure 1 for "CFTR mutation leads to intrinsic dysfunction in neutrophils from people with Cystic Fibrosis"

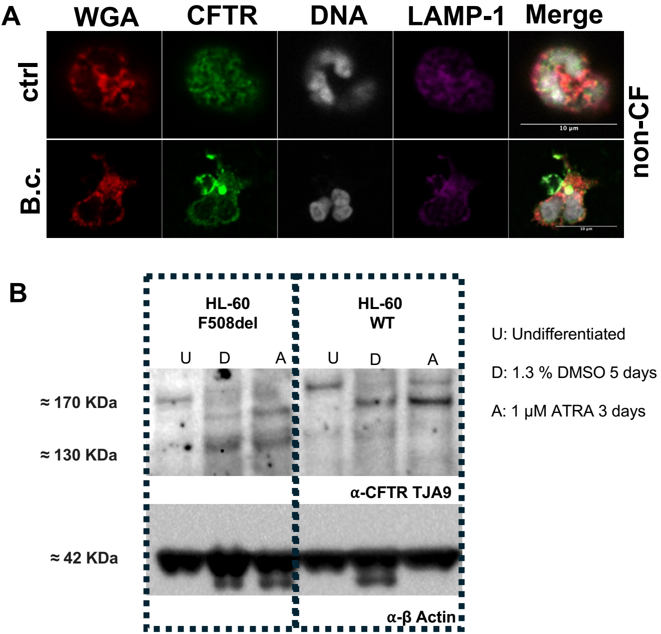


**Supplementary Figure 1. CFTR-deficient HL-60 human neutrophil-like cell line has a reduced CFTR mature expression. (A)** Non-CF neutrophils were seeded and infected with B. cenocepacia (MOI = 10) for 1 h, following for the fixation and staining with anti-CFTR (green), WGA (plasma membrane, red), DAPI (DNA, gray) and anti-LAMP1 (Lysosomes, magenta). **(B)** WT or F508del Undifferentiated HL-60 cells (U) or differentiated cells to neutrophil-like cells with 1.3% of DMSO (D) or with 1 μM retinoic acid (A) were lysed, and western blot was performed with anti-CFTR antibody [TJA9], the glycosylated band was detected at ≈ 170 KDa.
