## Supplemental Figure 2 for "CFTR mutation leads to intrinsic dysfunction in neutrophils from people with Cystic Fibrosis"

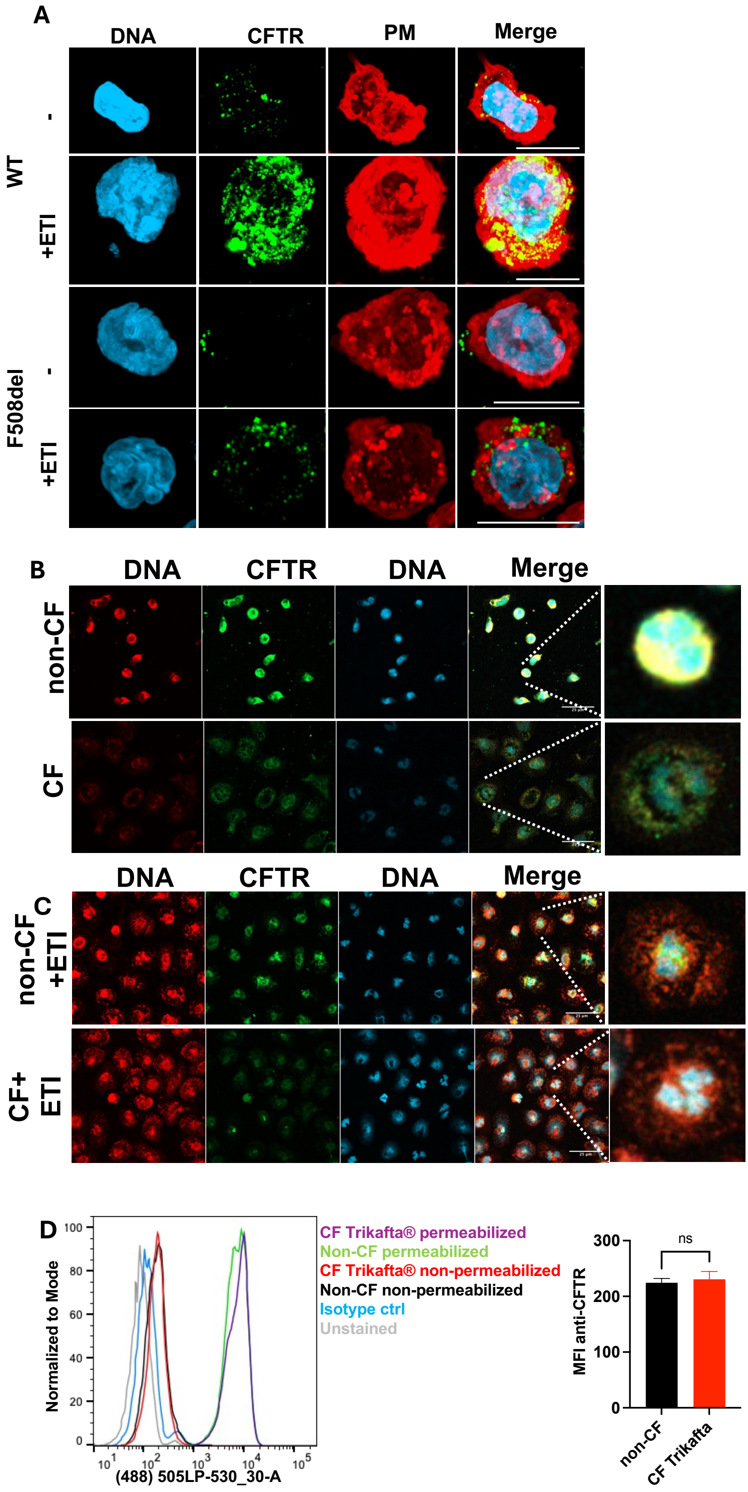


**Supplementary Figure 2. ETI increases the CFTR protein expression in F508del neutrophils.**

**(A)** WT and F508del HL-60 cells were treated with ETI and stained with anti-CFTR (green), WGA (plasma membrane, red) and DAPI (blue), F508del cells treated with ETI recovered the CFTR expression (n=3). **(B)** Non-CF or CF neutrophils were fixed, permeabilized and stained with anti-CFTR [CF3] (green), WGA (red) and DAPI (blue). **(C)** non-CF or CF neutrophils were treated in vitro with ETI and stained as mentioned before. Treating with ETI increased CFTR expression in neutrophils. (D) After staining with anti-CFTR, flow cytometry analyzed non-CF or CF neutrophils from pwCF treated with Trikafta®. Both groups revealed similar fluorescence in non-permeabilized or permeabilized conditions. The bar graph indicates similar CFTR expression levels (n=4, Welch's t-test).
