## Supplemental Figure 3 for "CFTR mutation leads to intrinsic dysfunction in neutrophils from people with Cystic Fibrosis"

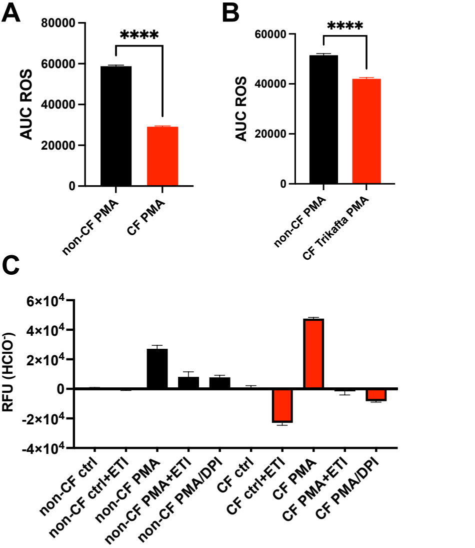


**Supplementary Figure 3. ETI restores the ROS production in CF neutrophils.**

The oxidative burst was measured in non-CF and CF neutrophils from pwCF before **(A)** and after Trikafta® treatment **(B)**, the ROS was detected with luminol after the stimulation with PMA, the Area Under the Curve were calculated and displayed as bar graphs (****=p>0.001, Welch's t test). **(C)** Hypochlorite production was measured in non-CF and CF neutrophils under Trikafta® treatment. The cells were stimulated with PMA and treated with ETI or DPI; aminophenyl fluorescein detected the hypochlorite. Relative Fluorescent Units (RFU) expressed the fluorescence.
